# RNA m^6^A Methylation Control Salt Response by Affecting Photosynthesis Capacity

**DOI:** 10.1101/2025.08.13.670251

**Authors:** Qinghua Yang, Shuang Wang, Jieying Si, Qiuying Pang, Aiqin Zhang

## Abstract

Soil salinization is a major abiotic stress constraining global plant growth. Epigenetic regulation particularly RNA modifications like N6-methyladenosine (m^6^A) is crucial for plant stress responses, the specific role of m^6^A in salt tolerance remains unclearly defined. Here, we show that Arabidopsis mutants deficient in m^6^A writers or readers exhibit heightened salt sensitivity at both growth and physiological levels. Global m^6^A modification levels increase following salt stress which results in a significant elevation of m^6^A signature throughout the Arabidopsis transcriptome. Spatiotemporal analysis through standardized read density of MeRIP-seq uncovered a significant salt-induced redistribution of m^6^A modification patterns. More than 80% m^6^A reads locate around the stop codon within 3’-untranslated regions and tend to increase under salt stress. A total of 9,986 peaks aligned into 8,667 coding gene showing significant changes in m^6^A modification levels. By associating with gene expression profiling, 840 salt-responsive genes display significant alterations in m^6^A enriched level and negative regulation pattern happens in most of genes. The salt responsive genes showing increased m^6^A methylation but significantly decreased expression are focused which representatively enriched in photosynthesis pathway, and this suppression due to mRNA decay mediated by m^6^A modification. Photosynthetic capacity and chloroplast apparatus are impaired in m^6^A-dependent manner under salt stress. These results clarify the biological functions of m^6^A modification in plant response to salinity and uncover the specific role of RNA metabolism based on dynamic change pattern of m^6^A modification to cope with salt stress.

## Introduction

Due to the sessile nature, plants have to confront with diverse abiotic stresses caused by climate change. Soil salinization is one of the worldwide adverse environmental factors that severely impedes plant growth, development and crop productivity (Yang and Guo, 2018). To carry out saline adaptations, plants have evolved sophisticated cellular and physiological mechanisms, which attribute to the global and rapid activation of genome-wide gene expression in plant responses to salt (Zelm et al., 2020). Approximately 30% of the transcriptome appears to change upon salt stress (Kreps et al., 2002). Among these, stimulus-specific genes undergo accurate regulation at chromatin, DNA and mRNA level to bring about appropriate responsiveness (Rasmussen et al., 2013; Singh and Roychoudhury, 2021). Epigenetic marks including DNA methylation, nucleosome remodeling and posttranslational modifications of the histones were well-characterized in plant salt tolerance (Zheng et al., 2016; Wang et al., 2021; Ueda and Seki, 2020; Yung et al., 2021). The achievement on the mapping of epitranscriptomic marks such as *N*^1^-methyladenosine (m^1^A), *N*^6^-methyladenosine (m^6^A) and 5-methylcytosine (m^5^C) gave impetus to investigate the potential role of RNA modifications in salt response (Dominissini et al., 2012; Luo et al., 2014; David et al., 2017; Liang et al., 2020; Gao et al., 2021). Recently emerging evidence has shown that the dynamics features of m^6^A RNA methylation have profound implications for post-transcriptional regulation of salt-responsive genes (Anderson et al., 2018; Arribas-Hernandez and Brodersen, 2020; Zheng et al., 2021; Hu et al., 2021).

m^6^A represents methylation at the N^6^-position of adenosine of messenger RNA (mRNA), which has been known as the most prevalent internal mRNA modification in all higher eukaryotes (Jia et al., 2013; Wang et al., 2014). m^6^A RNA methylation greatly determines the fate of mRNAs, lncRNAs and miRNAs, prompting differential mRNA degradation, stability, translation and then influencing gene expression (Visvanathan and Somasundaram, 2018). m^6^A methylome mapping shows the m^6^A marks predominantly locate in the 3_ untranslated regions (3_-UTRs) and near stop codons (Dominissini et al., 2012; Luo et al., 2014; 2020). It has been proposed that the distribution site of m^6^A is associated with alternative polyadenylation (APA) and subsequently affects mRNA stability and nuclear-to-cytoplasmic export (Yue et al., 2018; Song et al., 2021). In plants, m^6^A is cotranscriptionally installed by m^6^A methyltransferase complex. The core components serving as m^6^A writers include the methyltransferase-like proteins MTA and MTB, FKBP12 INTERACTING PROTEIN 37 KD (FIP37), KIAA1229/ VIRLIZER (VIR) and the E3 ubiquitin ligase HAKAI. All are essential for sculpting m^6^A mRNA modification (Zhong et al., 2008; Shen et al., 2016; Ruzicka et al., 2017). The functional disorder of any of these components results in plant growth and development defect (Liang et al., 2020). The m^6^A modification is reversible and the methyl groups of adenosines on mRNA can be removed by m^6^A demethylases that act as m^6^A erasers, such as ALKBH9B, ALKBH10B and SLALKBH2, which belong to ALKBH family proteins (Duan et al., 2017; Martinez-Perez et al., 2017; Zhou et al., 2019). These *N*^6^ methylated RNAs could be recognized by the reader proteins harboring the YTH (YT521-B homology) domain (Scutenaire et al., 2018). In *Arabidopsis*, the YTH family members EVOLUTIONARILY CONSERVED C-TERMINUS2 (ECT2), ECT3 and ECT4 specifically bind to m^6^A-modified mRNA and exert regulatory functions by mediating the stability of m^6^A-containing transcripts (Wei et al., 2018; Arribas-Hernández et al., 2020). Currently, given the functional demonstration in plants, m^6^A modification plays a vital role in multiple physiological processes, including embryo development, stem cell fate determination, leaf and root development, floral transition in *Arabidopsis*, sporogenesis and yield in rice, stress response in maize, apple and poplar, and fruit ripening in tomato (Yue et al., 2019; Liang et al., 2020; Yu et al., 2021;). In regard of stress responses, the global m^6^A level is sensitive to environmental changes, and the transcripts are subject to m^6^A modification during stress adaptation (Miao et al., 2020; Hu et al., 2021). Upon salt stress, recently study revealed that m^6^A is correlated with the mRNA stability of several salt response regulators and there was a link between m^6^A methylation and poly (A) site-dependent 3_-UTR length and mRNA stability (Hu et al., 2021). However, many important cellular pathways facilitating a proper physiological response to salt along with m^6^A status change remains ill defined. Owing to the robust and complicated molecular network of salt response regulation, the specific biological pathways processed in m^6^A-dependent manner under salt stress need to be ascertained.

In this study, by analyzing the Arabidopsis mutants of m^6^A writers and readers, we revealed the important role of m^6^A methylation in seedling establishment and plant growth under salt stress. Through m^6^A MeRIP-seq analysis of salt treated plants, we found that global m^6^A modification level increased following salt stress and most of m^6^A marks enriched in 3’-untranslated regions around the stop codon, showing tight association with the expression pattern of salt responsive gene. The genes involved in photosynthesis were overrepresented and underwent negative regulation mediated by m^6^A methylation, resulting in compromised photosynthetic capacity upon salt stress. Together, this study provides a novel perspective on understanding how dynamic m^6^A signatures regulate plant salt responsiveness and target specific physiological pathway by modulating transcription reprogramming.

## Materials and methods

### Plant materials and growth conditions

Col-0 accession of *Arabidopsis thaliana* was used as the wild-type (WT) plant. The T-DNA insertion lines of *mtb* (SALK_056904C), *fip37* (SALK-029377C), *vir* (SAIL-686-E12), *hakai* (SALK-148797C), *alkbh9b* (SALK-111811C), *alkbh10b-1* (SALK_004215C), *ect2-1* (SALK_002225C), *ect2-4* (SALK_146376C), *ect3-3* (SALK_134503C), *ect4-3* (SALK_112012C), *ca1* (SALK_106570C), *pnsb1* (SALK_206860C) and *lhcb2.4* (SALK_055354C) were obtained from the Arabidopsis Biological Resource Center (ABRC). The *ect* triple mutants were generated by genetic crossing. Seeds from WT and mutant lines were surface-sterilized and planted on half Murashige and Skoog (MS) medium plus 1% sucrose with 0.8% agar or in pots filled with uniformly mixed PINDSTRUP substrate, after stratification at 4 °C for 3 d, transferred to a controlled-environment growth chamber and cultivated at 22 °C with 50% relative humidity under a 16 h light/8 h dark (150 μM photons m^-2^ s^-1^) photoperiod. Salt stress was applied to three-week-old plants grown in soil treated with 150 mM NaCl for 72 h or directly added to the half MS medium. The primers used for genotyping of the mutants are listed in Supplementary Table S1.

### RNA extraction and quantitative reverse transcription-(qRT) PCR

Total RNA isolation from three-week-old plant leaves was carried out using TRIzol™ reagent (Thermo Fisher Scientific, USA) according to the manufacturer’s protocol. RNA integrity was verified by 2% agarose gel electrophoresis, followed by DNase I treatment (Takara, Japan) to remove DNA contamination. 2 μg RNA of each sample was used to cDNA synthesize by Maxima H Minus reverse transcriptase (Thermo Fisher Scientific, USA). The qRT-PCR analysis was performed using TB Green Premix Ex TaqTM (Takara, Japan) on an ABI 7500 Real-Time PCR System (Thermo Fisher Scientific, USA). Relative gene expression levels were quantified via 2^-ΔΔCt^ method, with *ACTIN2* serving as the endogenous reference gene. Three technical replications were conducted for each biological experiment. The primers used in the qRT-PCR analysis are shown in Supplementary Table S1.

### Phenotypic analysis of plant response to salt stress

Germination assay was performed on Col-0 and mutant lines sown on half MS agar plates supplemented with or without 150 mM NaCl. Each line contained 80 to 100 seeds for one plate. The germination rate based on radicle tip or green cotyledon emergence were recorded daily. 12-d-old seedlings grown on half MS medium were photographed and the pictures were processed with ImageJ software (download from http://imagej.nih.gov/ij/) to measure the root length. At least 30 seedlings were quantified per line, and each experiment was repeated three times.

### Dot blot assay

mRNA was isolated and purified using the Dynabeads^TM^ mRNA Purification Kit (Thermo Fisher Scientific, USA). The same amount of RNA was spotted on nitrocellulose membranes. Crosslink the membrane in UV-crosslinked for 1 min (1,200 microjoules [×100]) and block with 5% BSA. The blot was then probed with m^6^A antibody (ABclonal, China). Methylene blue staining was used as a loading control.

### mRNA stability assay

Wild-type and *vir* mutants were cultured on 1/2 MS medium for 7 days, then the seedlings were transferred to 1/2 MS liquid medium for growth and adaptation overnight, and the samples were collected after treatment with 10 μM actinomycin D for 0 h, 6 h, 12 h, and 24 h respectively. RT-qPCR was performed to determine mRNA levels, and statistical analysis was used to determine the significant difference between wild-type and mutant. The primers used in the study are listed in Supplementary Table S1.

### Chlorophyll fluorescence analysis

Chlorophyll fluorescence analysis was performed on three-week-old plants using a Fluor-Cam FC-800-O imaging fluorometer (Photon Systems Instruments) equipped with harmonically modulated actinic irradiance. Following dark adaptation, the minimal fluorescence *(F*_0_) was recorded, while the maximum fluorescence yield (*F*m) was determined under red actinic light (150 mmol m^-2^ s^-1^), from which the average quantum efficiencies of photosystem II (PSII) was derived. Key photosynthetic parameters including dark-adapted variable fluorescence *F*v and *F*m and Non-Photochemical Quenching (NPQ) were analyzed to evaluate the maximum PSII quantum yield.

### Histochemical detection of ROS accumulation

Visualization of superoxide and H_2_O_2_ in leaves was achieved through nitro blue tetrazolium (NBT) and 3,3’-diaminobenzidine (DAB) staining. Whole plants were vacuum-infiltrated with 1% (w/v) DAB in Tris-HAc (pH 6.5) for 24 h, and 0.5% NBT (w/v) in HEPES-K buffer (pH 7.0) under dark conditions at 28 °C for 4 h, then photographed after chlorophyll removal by 80% ethanol. At least five plants were processed per replicate for each genotype under control and salt stress.

### m^6^A MeRIP sequencing and data analysis

Total RNA of two-week-old seedlings grown on half MS medium supplemented with or without 150 mM NaCl was isolated and purified using TRIzol reagent (Thermo Fisher Scientific, USA). 0.5 mg plants were used for each RNA prep. RNA amount and purity of each sample was tested by NanoDrop ND-1000 (NanoDrop, USA). RNA integrity was assessed by gel electrophoresis and Bioanalyzer 2100 (Agilent, USA) with RIN number >7.0. Ribosomal RNA was removed from approximately 25 μg of total RNA representing a specific adipose type according to the protocol of the Epicentre Ribo-Zero Gold Kit (Illumina, USA). Following purification, the ribosomal-depleted RNA was randomly fragmented into ∼200 nt using Magnesium RNA Fragmentation Module (NEB, USA) under 86 _ for 7min. 0.5 μg fragmented RNA was used for RNA-seq analysis as Input, and 25 μg fragmented RNA was incubated for 2 h at 4 _ with m^6^A-specific antibody (Synaptic Systems, Germany) in IP buffer (50 mM Tris-HCl, 750 mM NaCl and 0.5% Igepal CA-630) with 0.5 μg_μL^-1^ BSA. The IP RNA was reverse-transcribed to create cDNA by SuperScript™ II Reverse Transcriptase (Thermo Fisher Scientific, USA), then used to generate U-labeled second-stranded DNAs with DNA polymerase I (NEB, USA), RNase H (NEB, USA) and dUTP Solution (Thermo Fisher Scientific, USA). Base A is added to the blunt ends of each strand for ligation to the indexed adapters. AMPureXP beads was used for size selection. After UDG enzyme (NEB, USA) treatment of the U-labeled second-stranded DNAs, the ligated products are amplified with PCR by the following conditions: initial denaturation at 95_ for 3 min, 8 cycles of denaturation at 98_ for 15 sec, annealing at 60_ for 15 sec, extension at 72_ for 30 sec, and final extension at 72_ for 5 min. MeRIP sequencing of the final cDNA library (300 ± 50 bp) was performed on Illumina Novaseq™ 6000 (LC-Bio Technology, China).

Raw paired-end reads of MeRIP and Input RNA sequencing were filtered and adapter sequences were trimmed out by fastp software (https://github.com/OpenGene/fastp). Cleaned reads were aligned to the *Arabidopsis* reference genome (v48) using HISAT2 software (http://daehwankimlab.github.io/hisat2). Mapped reads of IP libraries were provided for R package exomePeak (https://bioconductor.org/packages/exomePeak) to identify m^6^A peaks with the input as background upon bigwig format that can be adapted for visualization on the IGV platform (http://www.igv.org). MEME (http://meme-suite.org) and HOMER (http://homer.ucsd.edu/homer/motif) were used for de novo and known motif finding followed by localization of the motif with respect to peak summit. Called peaks were annotated by R package ChIPseeker (https://bioconductor.org/packages/ChIPseeker). mRNA expression analysis of Input libraries was performed using StringTie (https://ccb.jhu.edu/software/stringtie) based on FPKM calculation. The differentially expressed mRNAs were selected with log2 (fold change) > 1 or log2 (fold change) < -1 and *p* value < 0.05 by R package edgeR (https://bioconductor.org/packages/edgeR). Three biological replicates were taken for each sample under control and salt stress.

### RNA sequencing

2 μg of total RNA were used for mRNA purification and cDNA synthesis. The cDNA (100∼200 bp) samples were purified with AMPure XP system (Beckman Coulter, USA). PCR Enriched cDNAs were used to create the final cDNA library and library quality was assessed with the Agilent Bioanalyzer 2100 system (Agilent Technologies, USA). The high-performance sequencing was run on Illumina HiSeq 2500 platform. Clean reads were mapped to the Arabidopsis TAIR10 reference genome and assembly were performed with the software tools HISAT2 and StringTie. Differential expression analysis was performed using DESeq2 package. Genes with an FDR-adjusted fold change ≥2 and p ≤ 0.05 were considered to be differentially expressed between two samples. AgriGO and TopGO software was used for gene ontology (GO) functional classification and enrichment. Three biological replicates were taken for each genotype under control and salt stress conditions.

### m^6^A-IP-qPCR

m^6^A-IP-qPCR was performed to validate the m^6^A enrichment at specific loci of selected mRNA. 14-d-old WT seedlings were used for the IP assays as mentioned above. The m^6^A-containing fragments were immunoprecipitated with preblocked Protein A Dynabeads (Thermo Fisher Scientific, USA). The mRNA fragments treated with beads only but without IP was taken as the Input control. All the samples were subjected to reverse transcription and qPCR assays after ethanol precipitation. m^6^A enrichment was estimated as the fraction of IP mRNA relative to Input (% Input). Three biological replicates were performed. Sequences for the primers used for IP-qPCR are listed in Supplementary Table S1.

### Transmission electron microscopy observation

The plant samples were primarily fixed in 2.5% (w/v) glutaraldehyde in 0.1 M phosphate buffer (pH 7.4) for 24 h at 4°C, followed by secondary fixation with 1% osmium tetroxide (OsO4) in the same buffer for 2 h. Subsequently, the samples underwent graded ethanol dehydration (30%, 50%, 70%, 85%, 95%, and 100%) followed by acetone transition. The dehydrated specimens were infiltrated with Spurr’s epoxy resin through a graded series (25%, 50%, 75%, and 100%) and polymerized at 60°C for 48 h. Ultra-thin sections (70-90 nm) were obtained using a diamond knife-equipped ultramicrotome (Leica EM UC7). The sections were double-stained with 2% uranyl acetate in 50% ethanol and Reynolds’ lead citrate, then examined under a transmission electron microscope (HITACHI HT7700) operated at 80 kV accelerating voltage.

### Accession numbers

Sequence data in this study can be found in TAIR with the following accession numbers: *mtb* (SALK_056904C), *alkbh10b-1* (SALK_004215C), *ect2-1* (SALK_002225C), *ect2-4* (SALK_146376C), *ect3-3* (SALK_134503C), *ect4-3* (SALK_112012C), *ca1* (SALK_106570C), *pnsb1* (SALK_206860C) and *lhcb2.4 (*SALK_055354C).

### Statistical analysis

Student’s *t-*test was used to assess the statistically significant differences between wild type and mutants under salt stress, respectively. Multi-factor ANOVA analysis for genotype-environment interaction (G×E) was performed using Tukey’s multiple comparison test in the RStudio 1.2.1335 (https://www.rstudio.com/).

## Results

### The change of m^6^A level affect the response to salt stress

To elucidate the involvement of m^6^A regulatory components in response to salt stress, we systematically analyzed the temporal expression profiles of the genes encoding key m^6^A-modulating factors including m^6^A writers, erasers and readers using qRT-PCR. Wild-type (Col-0) seedlings subjected to 200 mM NaCl treatment over a 24-hour time course (0, 3, 6, 12, and 24 h) demonstrated significant transcriptional reprogramming of m^6^A-associated genes. The core writer complex components *MTA*, *MTB*, *VIR*, *FIP37* and *HAKAI* exhibited progressive upregulation (1.1- to 7.69-fold) upon salt exposure (Fig 1A). Parallel analysis revealed enhanced expression of eraser genes *ALKBH9B* and *ALKBH10B* (Supplementary Fig S1). Notably, most of the readers, such as *ECT2*, *ECT3* and *ECT4* displayed marked induction (more than 6-fold) after salt stress (Fig 1B). To corroborate these transcriptional changes, we quantified global m^6^A levels via dot blot immunoassay. A significant 15% elevation (*P* < 0.01) in global m^6^A modification was detected in stressed seedlings compared to the controls (Fig 1C, D). These coordinated transcriptional activation events and concomitant m^6^A hypermethylation suggest that dynamic epitranscriptomic reprogramming through m^6^A modification gets involved in plant salt response.

**Fig. 1.**
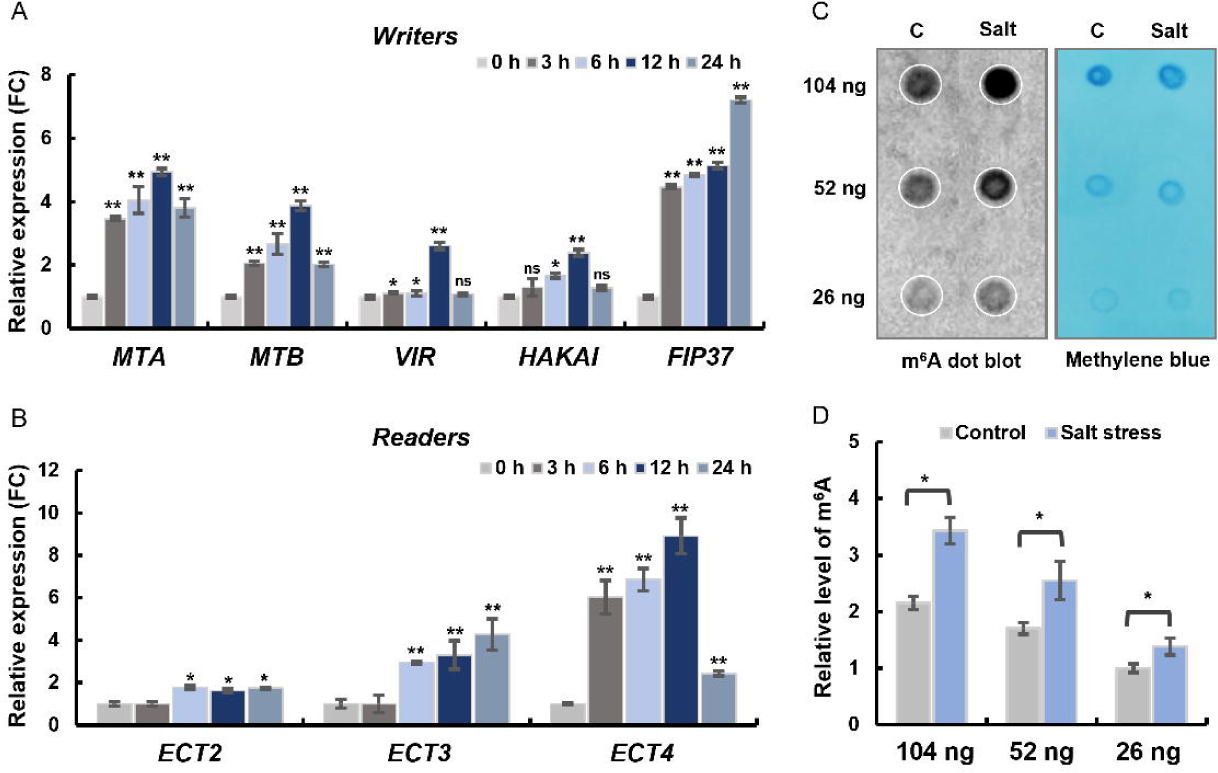
The effect of salt stress on m6A modification and the expression of its related genes. **A)** RT-qPCR analysis for the temporal expression profiles of the genes encoding key m6A writers in Col-0 seedlings during 24 hours of salt treatment. **B)** Expression levels of the key genes encoding m6A reader in Col-0 seedlings during salt treatment. **C)** Dot blot assay showing the levels of m6A modification in poly(A+) mRNA isolated from Col-0 seedlings after 7 days of salt treatment. Methylene blue staining represents the loading control. **D)** Quantification of m6A levels by dot density. 2 µL of 104, 52 and 26 ng poly(A+) mRNA from the same replicate are blotted onto the nylon membrane. n = 3 biological replicates. Statistically significant differences compared with the control condition were assessed using paired Student’s t-test: *\*p* < 0.05, ** *p* < 0.01, ns, not significant.

To further elucidate the functional significance of m^6^A methylation in salt stress response, we used the mutant of the key member of m^6^A writers to analyze the salt-responsive phenotypes. Previous report has been shown m^6^A level significantly decreased in the mutants of all writers. In view of the interference of the growth defect in these mutants, we first used *mtb* plants to analyze the involvement of m^6^A in salt response. According to the literature, *mtb* is loss-of-function mutants of the gene (Wu et al., 2020). The primary root length of *mtb* seedlings was strongly inhibited under salt stress when compared to Col-0 (Fig 2A, B), revealing hypersensitivity to salt stress.

**Fig. 2.**
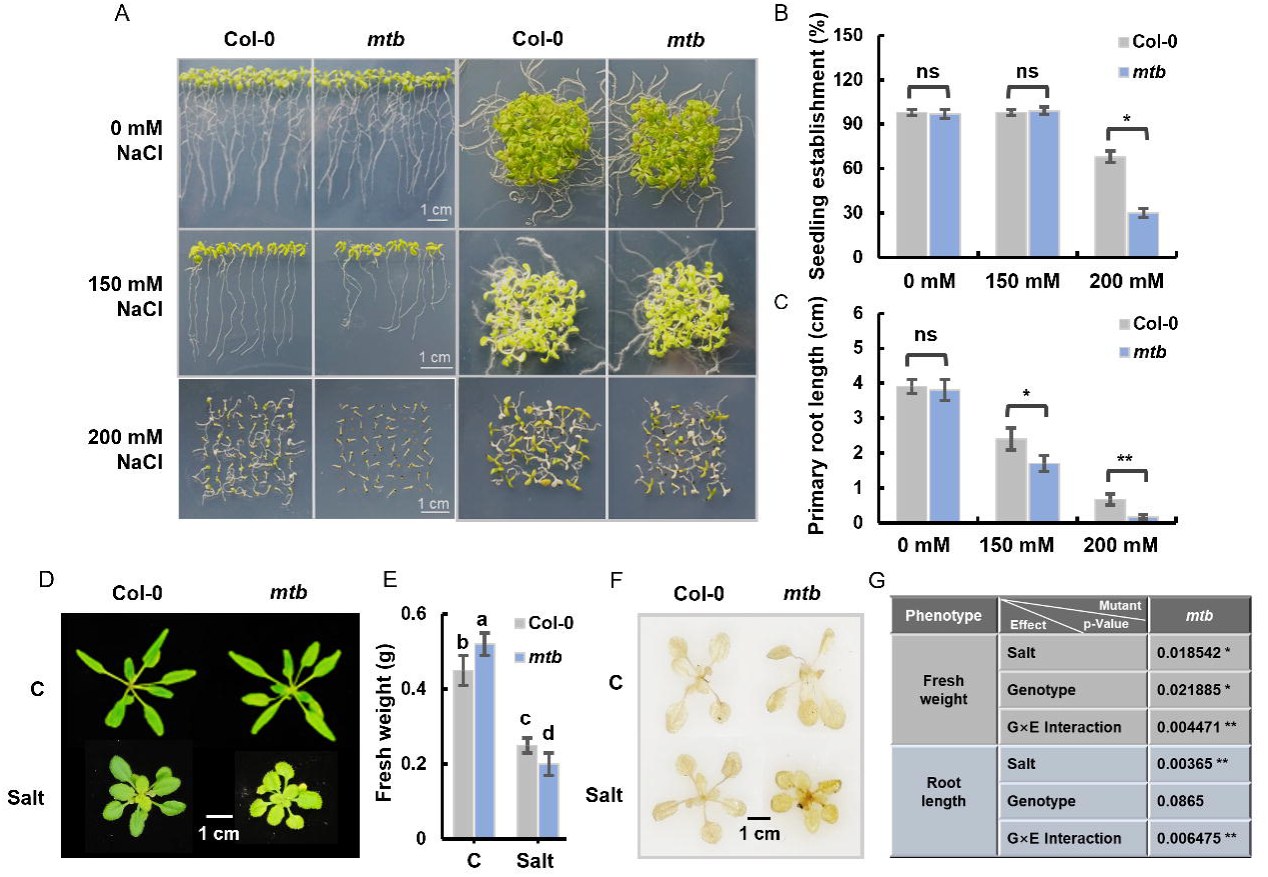
The growth phenotype and physiological response of m6A writer mutant *mtb* under salt stress. **A)** Phenotypic determination and quantification of *mtb* mutants after 14 days of 150 mM and 200 mM NaCl treatment, measuring seedling establishment rate(%) **B)** and root length **C)** compared to Col-0 plants. Scale bar= 1 cm. **D)** The phenotype of rosette growth and **E)** fresh weight of Col-0 and *mtb* plants grown in pots for 3 weeks and then subjected to 7 days of salt treatment. Different letters represent significant difference between treated conditions and genotypes (Duncan’s test,p < 0.05). **F)** 3.3’-diaminobenzidine (DAB) staining in Col-0 and *mtb* mutants before and after salt treatment. Scale bar = 1 cm. **G)** Factor analysis of genotype by salt (GxE) interaction on *mtb* mutant, the *p* values of effects of salt, genotype and G×E interaction on growth phenotypes. Significance is indicated by asterisk: *\*\*p* < .01; *\*p* < *.05;p* > .05, not significant.

The emergence of cotyledon and seeding establishment rate of *mtb* displayed a reduction compared with wild type in the presence of 200 mM NaCl (Fig 2C). Under normal conditions, wild-type and mutant plants showed a healthy and green leaf morphology (Fig 2D). Salt treatment significantly increased the sensitivity of *mtb* to salt stress, accompanied by a decrease in fresh weight (Fig 2E) and an increase in ROS accumulation (Fig 2F). Through factor analysis of genotype-environment interactions, we demonstrated that the biomass accumulation and root development of *mtb* mutants under salt stress exhibit significant G×E effects (*p* < 0.01). Then the other writer mutants were all included to test the salt sensitivity. Comparative phenotypic analysis revealed *vir*, *fip37* and *hakai-2* mutants exhibited the most pronounced stress susceptibility as quantified by fresh weight reduction and enhanced ROS accumulation (Supplementary Fig S2). These coordinated changes suggest that m^6^A writer deficiency compromises cellular capacity to mitigate salt stress. The salt responsiveness was also assessed in the mutants of m^6^A erasers and readers. There was no significant phenotype change observed in *alkbh9b*, *alkbh10b-1*and *alkbh9b/10b* (Supplementary Fig S3), whereas *ect* mutants slightly sensitive to a relatively severe salt stress (Supplementary Fig S4A). More seedlings with albino phenotype appeared in *ect2*, *ect3* and *ect4* (SSupplementary Fig S4B). The triple mutant of *ect2/3/4* exhibited longer root length and higher fresh weight under control conditions (Fig 3A, B). Through factorial analysis of genotype-environment interactions, we demonstrated a significant G×E effect (*p* < 0.001) on both biomass accumulation and root development in *ect2/3/4* mutants under salt stress. Notably, the salt hypersensitivity was specifically attributed to the genetic ablation of *ECT2/3/4* rather than pleiotropic growth differences, as evidenced by comparable growth parameters between mutants and wild-type controls under normal conditions (Fig 3F). Taken together, these results indicate that the dysfunction on m^6^A modification as well as recognition give rise to bereft of plant salt response.

**Fig. 3.**
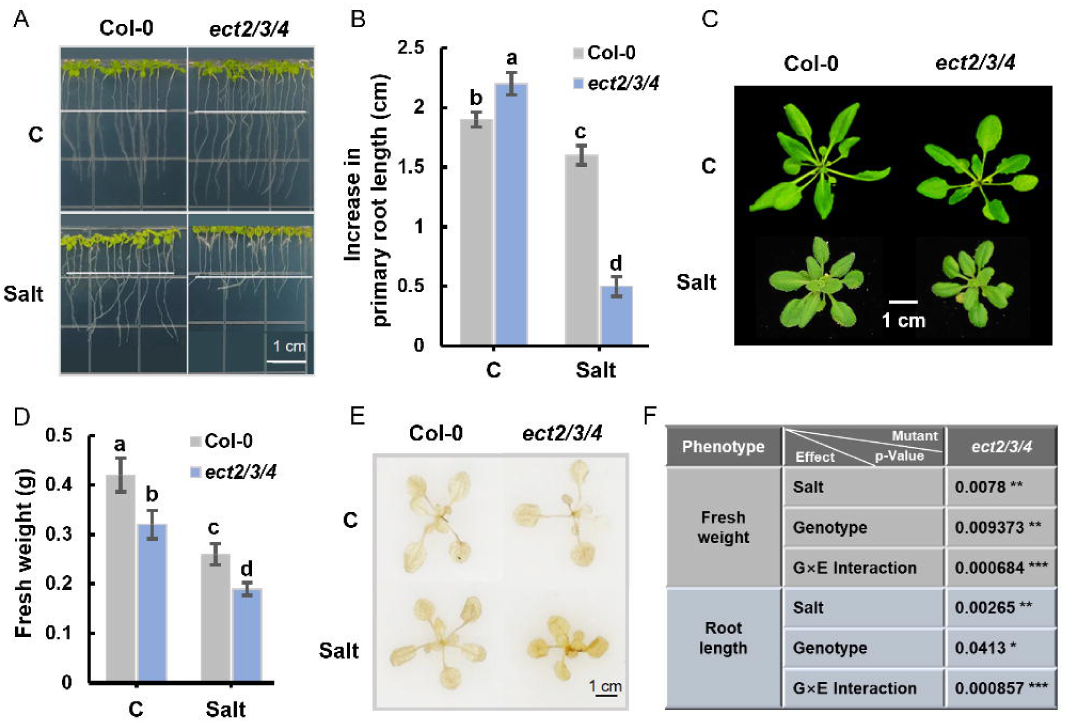
Effect of salt stress on the growth of m6A reader mutant *ect2/3/4.***A)** The seedling growth phenotype and **B)** determination of increase in primary root length of *ect2/3/4* mutants under 150 mM NaCl treatment for 7 days. **C)** The phenotype of rosette growth and **D)** fresh weight of Col-0 and *ect2/3/4* plants grown in pots for 3 weeks and then subjected to 7 days of salt treatment. Different letters represent significant difference between treated conditions and genotypes (Duncan’s test, *p* < 0.05). **E)** DAB staining in Col-0 and *ect2/3/4* mutants before and after salt treatment. Scale bar= 1 cm. **F)** Factor analysis of G×E interaction on *ect2/3/4* mutant, the *p* values of effects of salt, genotype and G×E interaction on growth phenotypes. Significance is indicated by asterisk: *\*\*\*p* < .001; *\*\*p* < .01; *\*p* < .05; *p* > .05, not significant.

### Features of m^6^A mapping response to salt

To obtain the transcriptome-wide m^6^A map in response to salt stress, m^6^A-immunoprecipitation sequencing was conducted using the m^6^A-targeted antibody coupled with matched input RNA sequencing for the seedlings with or without salt treatment (Supplementary Table S2). For each set, the m^6^A peaks were highly consistently among three biological replicates. These well-recurring peaks with high confidence were used for further analysis. A total of 7,489 m^6^A peaks within 3,970 coding gene transcripts in control plants, and 6,094 m^6^A peaks within 3,970 coding gene transcripts in salt treated plants were identified (Supplementary Table S2). Among them, 4,317 m^6^A peaks were detected within both group (*p* < 1e-5, Chi-squared test) (Fig. 1b-c, Supplementary Table S4). Comparing to control plants, global profiling revealed a significant elevation in m^6^A methylation levels throughout the *Arabidopsis* transcriptome following salt stress treatment (Fig 4A). The read distribution analysis showed that the reads from m^6^A containing transcripts (more than 80%) were predominantly located around the stop codon and within 3′-untranslated region (3′-UTR) in both growth conditions, while less present in 5′-UTR and coding sequences, and more peaks prone to enrich in 3′-UTR under salt stress (Fig. 4b-d), additionally demonstrating a global change on the landscape of m^6^A deposition upon salt stress.

**Fig. 4.**
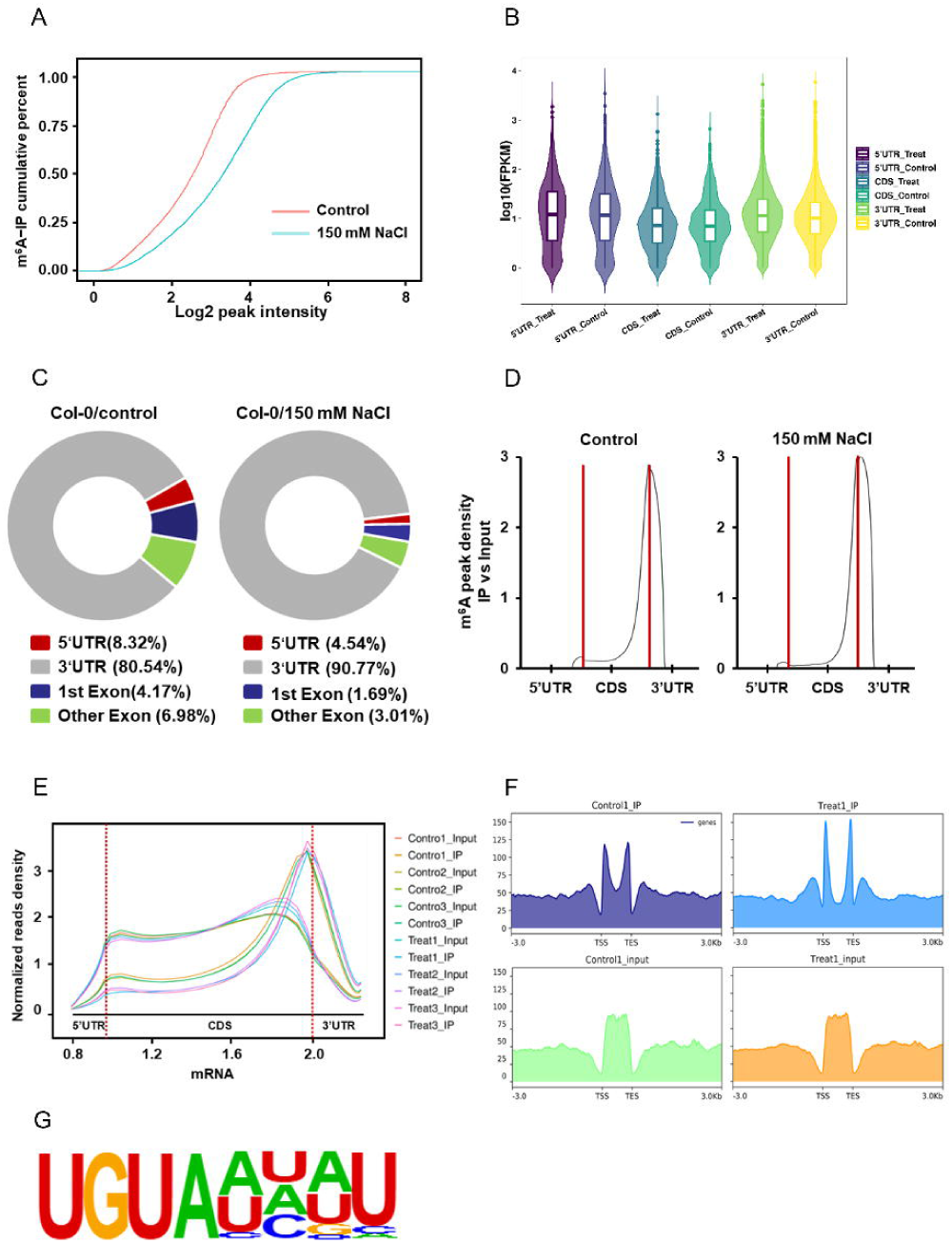
Genome-wide distribution of m6A in wild type Col-0 under salt stress. **A)** CDF plot of log converted fold changes of genes with control unique peaks (red line) or 150 mM NaCl treated unique peaks (blue line). Estimated (DESeq2) fold changes of genes in control and NaCl treated samples are plotted on the x axis. **B)** The enrichment of m6A peaks in the wild type Col-0 under non-stress and salt stress conditions. **C)** The distribution of m6A peaks in genes regions identified at Col-0 before and after salt treatment. **D)** The density of normalized m6A peaks along the mRNA transcripts before and after salt treatment in 5’UTR, CDS, and 3’UTR regions. **E)** Metaplots of m6A enrichments along normalized m6A peaks of Input and IP groups before and after salt treatment. **F)** The m6A signal of IP samples in the control and salt treated group was specifically enriched at the transcription start site (TSS) and transcription termination site (TES). **G)** RNA motifs identified by HOMER to be enriched in the indicated m6A peak regions. Three biological replicates are shown.

Spatiotemporal analysis through standardized read density uncovered a significant salt-induced redistribution of m^6^A modification patterns. While control conditions showed relatively uniform modification across UTR regions, salt-treated samples exhibited a distinct spatial shift characterized by reduced signal intensity in 5_-UTR and enhanced modification density surrounding stop codon region and throughout 3_-UTR (Fig 4E). Through in-depth analysis of different functional regions of the gene, it was observed that the m^6^A signal in the control IP samples was specifically enriched at the transcription start site (TSS) and transcription termination site (TES), and the signal intensity in this area was significantly increased by salt treatment (Fig 4F). Sequence motif analysis of m^6^A-modified transcripts obtained from all m^6^A MeRIP-seq data revealed that one conserved sequence is UGUAH (H = A, C, or U) (Fig 4G), which are plant-specific motifs (Shen et al., 2016).

### m^6^A affects mRNA stability of photosynthesis related genes

To further understand the involvement of m^6^A methylation in the salt stress response, we compared all m^6^A peaks detected under normal and salt stress conditions. A total of 9,986 peaks aligned into 8,667 coding gene showing significant changes in m^6^A modification levels were identified, and the distribution of these differential m^6^A peaks within gene regions was examined, revealing that the differential m^6^A peaks are primarily located in the 3’ UTR (Fig 5A). To further investigate the relationship between changes in m^6^A modification under salt stress and the regulatory activities of plant transcriptional responses to salt, an association analysis of MeRIP-seq and RNA-seq was conducted between salt-treated and untreated wild-types. Genome-wide analysis identified 8,667 genes exhibiting significant alterations in m^6^A modification levels under salt stress. By correlating these changes with the 3,135 differentially expressed genes under salt stress (Supplementary Table S3), we found that 840 salt-responsive genes displayed significant alterations in m^6^A modification (Fig 5B). Further analysis indicates that changes in m^6^A modification are associated with multiple modes of gene expression regulation (Supplementary Table S5). Specifically, we observed the following patterns: (1) Among genes with upregulated m^6^A modifications, 216 exhibited positive correlation between expression levels and modification abundance, while 86 showed negative correlation; (2) Among genes with downregulated m^6^A modifications, 374 displayed significantly increased expression, whereas 277 showed suppressed expression (Fig 5C, D). These results indicate that changes in m^6^A modifications exert multifaced regulatory effects on the expression of salt-responsive genes, though predominantly negative regulation was observed. Given existing evidence that m^6^A modification recruit recognition proteins (e.g., YTHDF2) to mediate mRNA degradation and reduce its stability (Wang et al., 2014). we focused on 86 salt-responsive genes showing increased m^6^A methylation but significantly decreased expression. We hypothesize that elevated m^6^A levels in these genes recruit m^6^A readers, accelerating mRNA degradation. Functional enrichment analysis revealed significant overrepresentation in photosynthesis pathway, particularly components of photosystem II (*OHP1*, *LHCB2.4*), Calvin cycle enzymes (*PRK*, *CA1*), chloroplast ATP synthase (*ATPD*), and NDH complex subunits (*PNSB1*) (Fig 5E). IGV visualization further confirmed pronounced enrichment of m^6^A modification peaks near stop codons under salt stress (Fig 5F). This prompted focused analysis of key photosynthesis-related genes, including light-harvesting complex-like protein *OHP1* (*OHP1*), phosphoribulokinase (*PRK*), ATP synthase subunit delta (*ATPD*), photosynthetic NDH subunit of subcomplex B 1 (*PNSB1*), beta carbonic anhydrase 1 (*CA1*), and Chlorophyll a-b binding protein 2.4 (*LHCB2.4*). Independent m^6^A IP-qPCR confirmed elevated m^6^A levels in these genes under salt stress in wild-type plants (Fig 6A-F), consistent with MeRIP-seq data showing substantial enrichment of m^6^A marks in their 3_-UTRs. RT-qPCR analysis revealed concomitant downregulation of these genes, suggesting m^6^A-mediated suppression (Fig 6G-L).

**Fig. 5.**
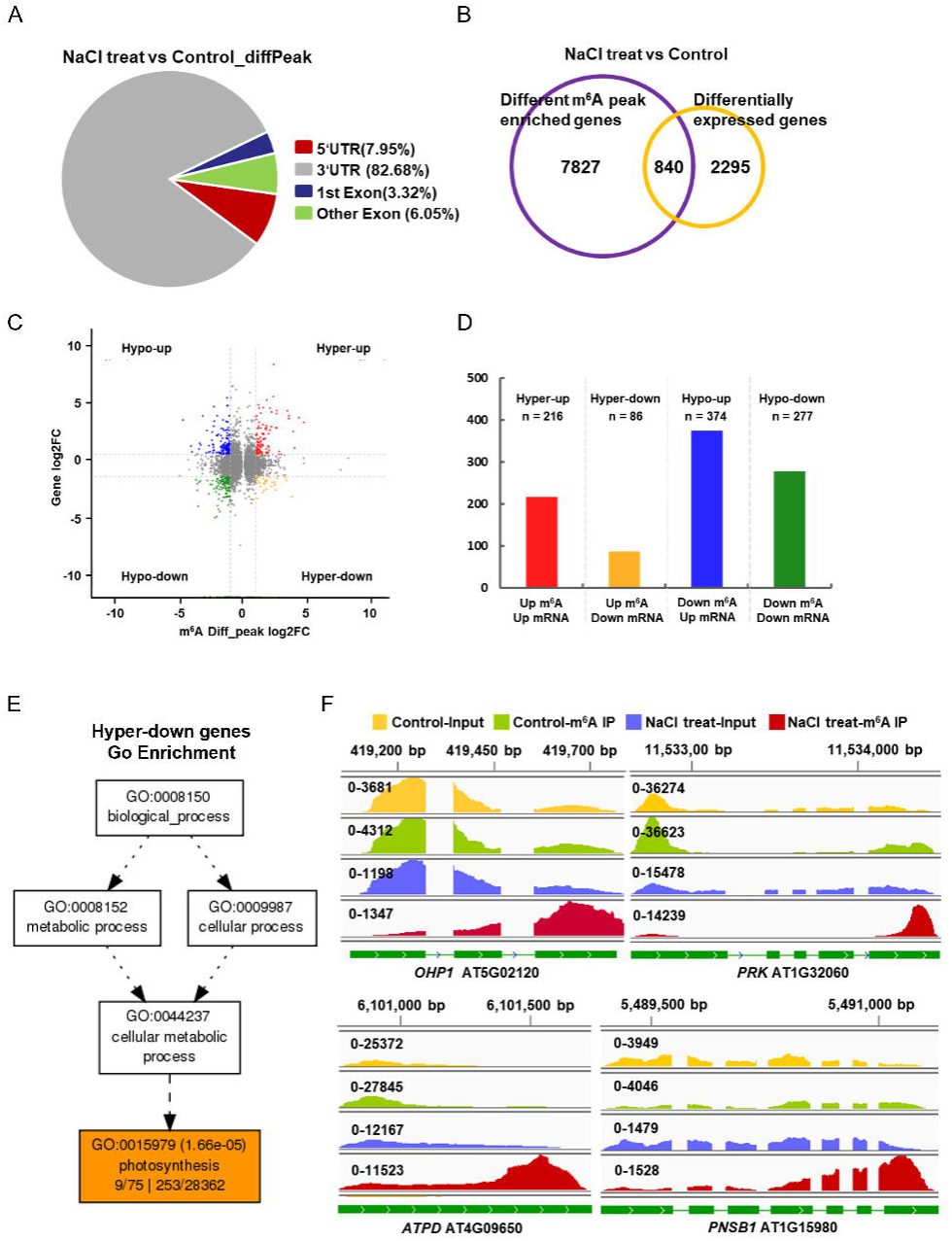
The dynamic change of m6A modification correlates with the different levels of gene expression upon salt stress. **A)** Distribution of differential m6A peaks in gene regions before and after salt treatment. **B)** The overlap between genes with significant changes in m6A modification and differentially expressed genes under salt stress. **C, D) The** regulatory mode of m6A modification changes on the gene expression pattern response to salt stress. Red dots (Hyper-up), hypermethylation and upregulated genes; yellow dots (Hyper-down), hypermethylation and downregulated genes; blue dots (Hypo-up), hypomethylation and upregulated genes; green dots (Hypo-down), hypomethylation and downregulated genes. **E)** Gene Ontology (GO) term enrichment of DEGs with m6A hypermethylation and downregulated genes. **F)** Representative Integrative Genomics Viewer (IGV) plot showing m6A peaks detected in photosynthetic genes under salt stress.

**Fig. 6.**
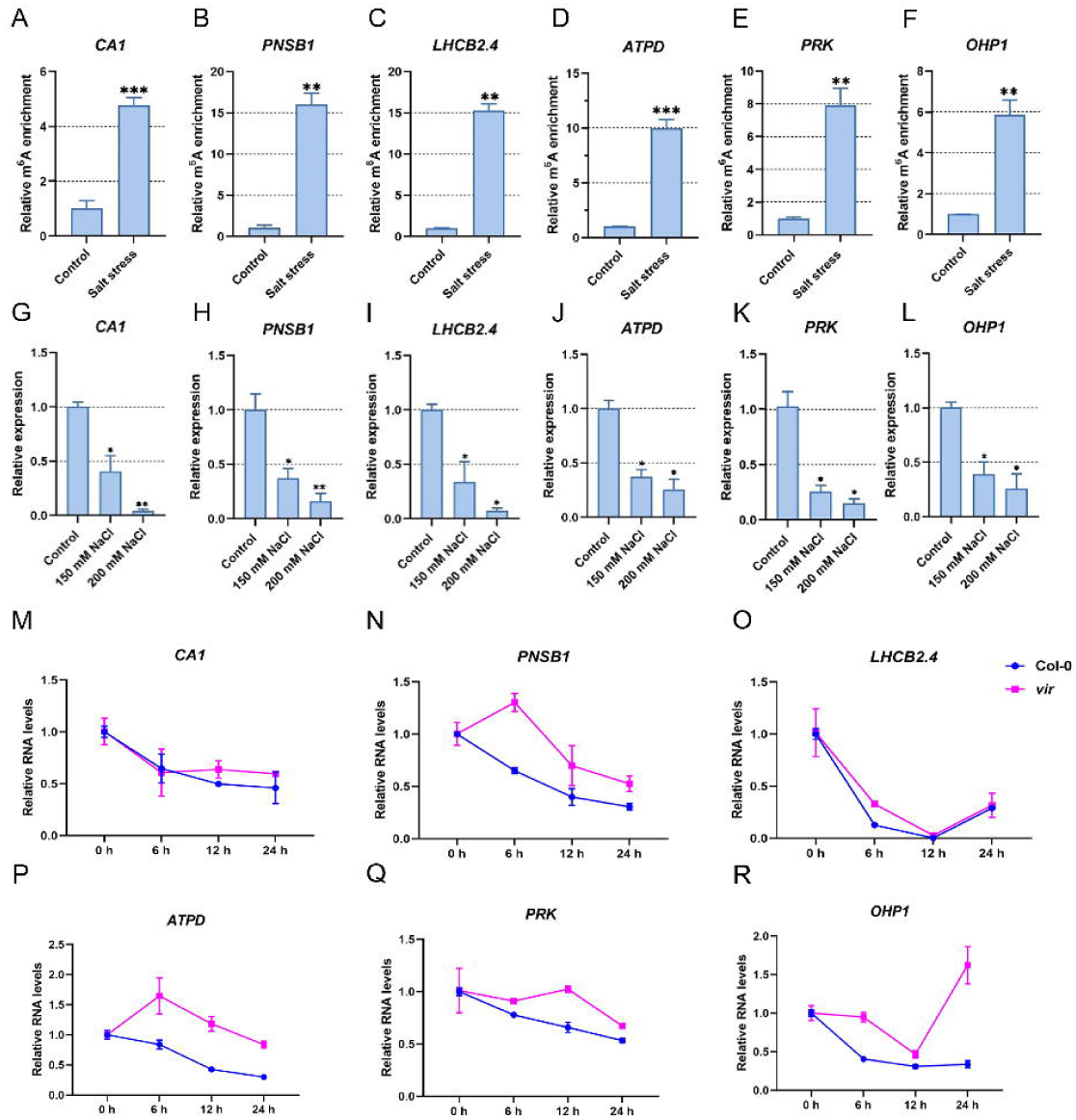
Salt stress affects the expression of photosynthetic genes depends on m6A modification. **A)** m6A-IP-qPCR analysis showed the m6A enrichment in *CAI,* **B)** *PNSBJ,* **C)** *LHCB2.4,* **D)** *ATPD,* **E)** *PRK* and **F)** *OHPJ* after salt treatment. **G-L)** The expression levels of photosynthetic genes after salt treatment detected by qRT-PCR analysis. Statistically significant differences compared with the control condition were assessed using paired Student’s t-test: *\*p* < 0.05, ** *p* < 0.01. **M-R)** mRNA stability assay for photosynthetic genes in wild type Col-0 and *vir* mutants after 3-DA treatment. Seedlings were treated with actinomycin D for 0, 6, 12 and 24 h. n = 3 biological replicates.

Given the established role of m^6^A in modulating RNA stability, we hypothesized that increased methylation accelerates transcript decay. We performed mRNA decay assays using *vir* mutant treated with Actinomycin D to inhibit transcription. Transcripts of *CA1*, PNSB1, LHCB2.4, ATPD, PRK, and OHP1 exhibited accelerated degradation in wild-type (Col-0) compared to *vir* mutants (Fig 6M-R). These results demonstrate that 1) VIR-dependent m^6^A methylation promotes mRNA decay of photosynthesis-related genes, and 2) salt-induced m^6^A hypermethylation suppresses their expression through enhanced transcript degradation.

### m^6^A modification maintains plant photosynthetic capacity under salt stress

To investigate the salt response of m^6^A-regulated photosynthetic genes, we obtained T-DNA insertion mutants (*ca1*, *pnsb1*, *lhcb2.4*), of which *ca1* and *pnsb1* are identified as loss-of-function mutants, while *pnsb1* is a functional deficiency mutant (Supplementary Fig S5). We then assessed salt sensitivity by comparing growth phenotypes of three-week-old mutants and wild-type (WT) plants under salt stress. Under control conditions, all genotypes displayed robust growth with vibrant green leaves, though mutants showed slightly reduced fresh weight, presenting a mild growth-deficient phenotype. Following salt treatment, WT and *ca1* leaves darkened, whereas *pnsb1* and *lhcb2.4* mutants developed yellow-green leaves with abnormal downward curling (Fig 7A). Salt stress significantly reduced fresh weight across all genotypes, but WT maintained higher biomass than mutants (Fig 7B). These results demonstrate that impaired function of CA1, PNSB1, or LHCB2.4 confers salt sensitivity. Genotype × environment interaction analysis confirmed these phenotypes resulted from gene deletion rather than inherent growth defects (Supplementary Fig S6).

**Fig. 7.**
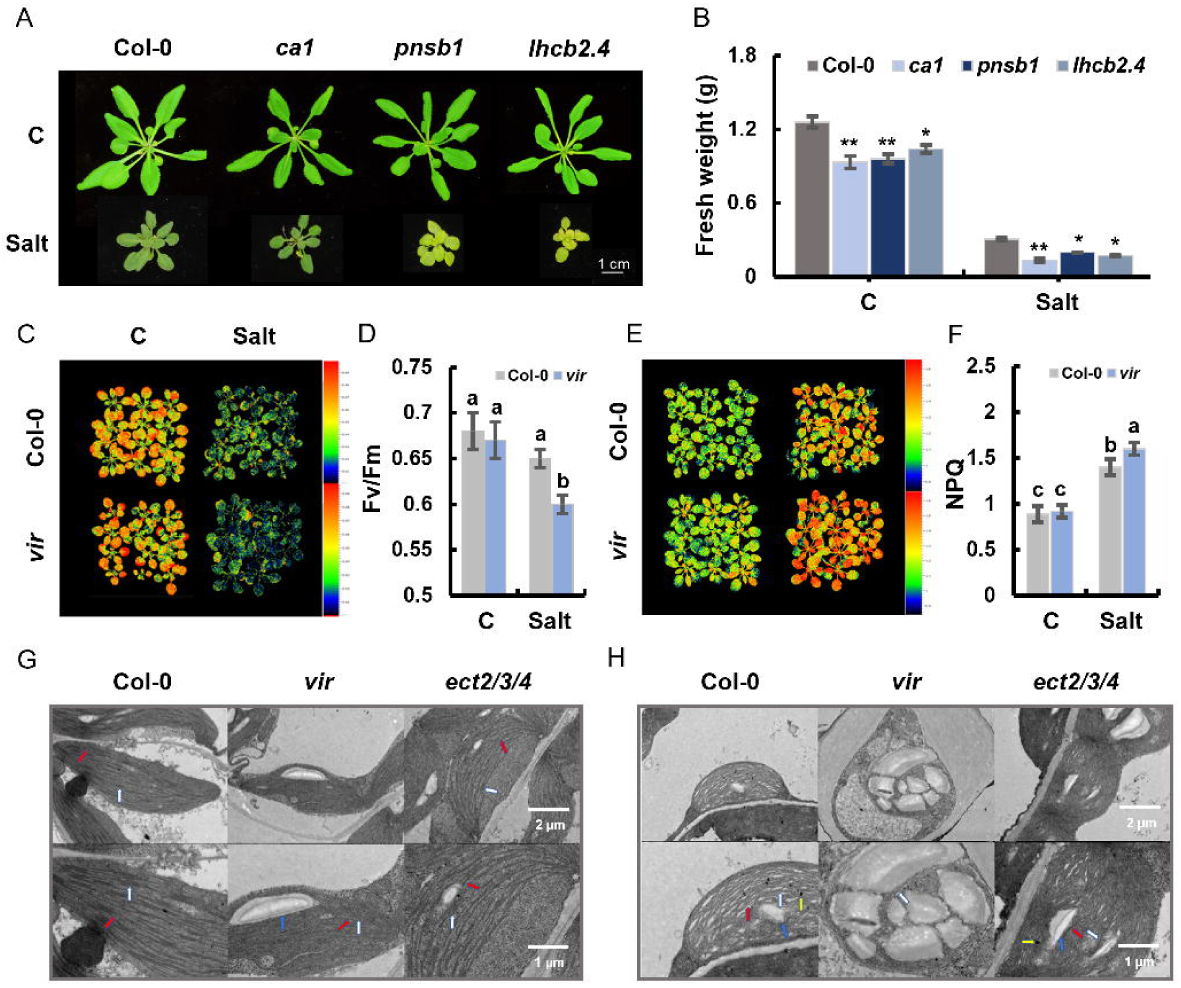
Salt induced photosynthetic deficiency in m6A-related mutant *vir* and *ect2/3/4.* **A)** The rosette growth and **B)** fresh weight of Col-0 and photosynthetic gene-associated mutants grown in pots for 3 weeks and then subjected to 7 days of salt treatment. **C-D)** Fv/Fm ratios showing the maximum quantum efficiency of PSII in Col-0 and *vir* plants before and after salt treatment. **E-F)** NPQ refers to Non-photochemical quenching. Values were counted by using Photon Systems Instruments, PSI system. Different letters represent significant difference between treated conditions and genotypes (Duncan’s test, p < 0.05). **G)** TEM imaging of chloroplasts from Col-0, *vir* and *ect2/3/4* showing typical internal chloroplast structures under non-stress condition. White arrows show the complete thylakoid system stacked as grana, red arrows show grana connected by stroma lamellae, the blue arrows show the starch granules ( Scale bar = 2 µm) **H)** TEM imaging of chloroplasts from Col-0, *vir* and *ect2/3/4* showing typical internal chloroplast structures after 14 days of salt treatment. White arrows show the dissolved thylakoid system, red arrows show the disintegrated stroma lamellae, yellow arrows show the plastid globules, and blue arrows show the starch granules( Scale bar= 2 µm.)

**Fig. 8.**
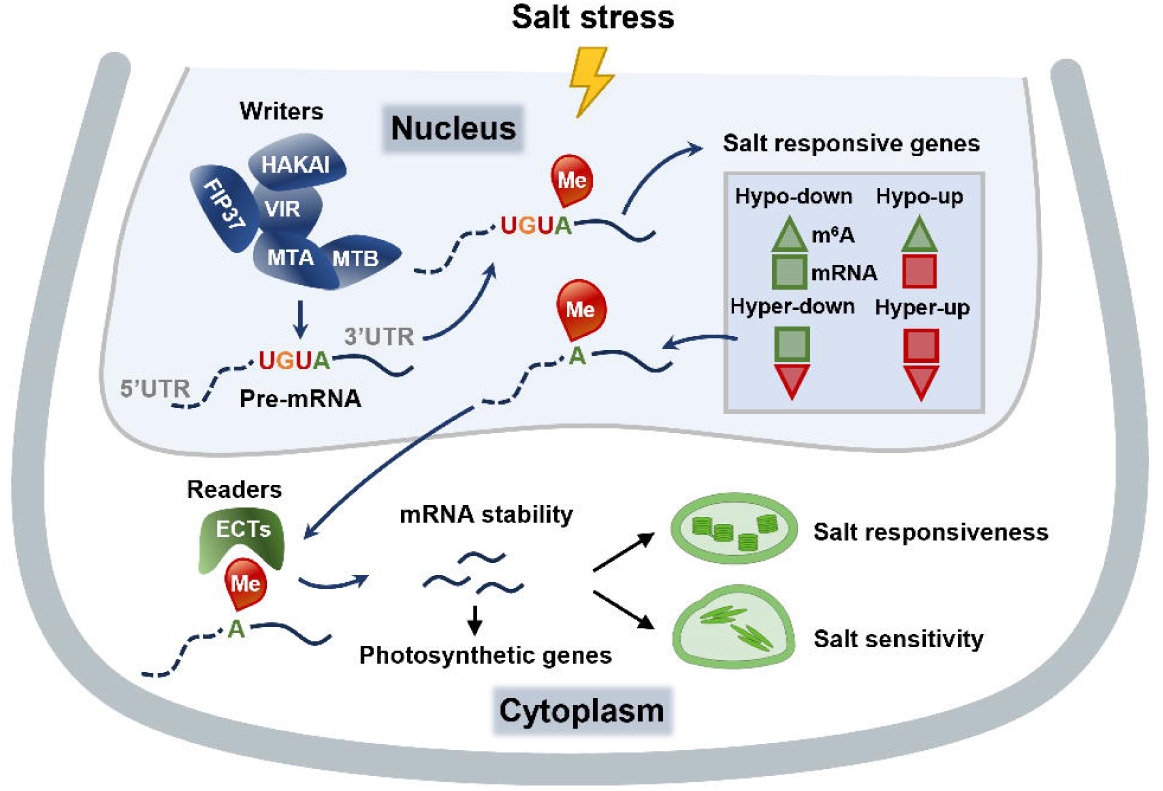
A model depicting the function of global m6A methylation in modulating plant salt response. Transcripts of m6A writers are induced by salt stress, prompts a significant increase of m6A level across Arabidopsis genome, which boosts the plant transcriptional responsiveness upon salt stress. The global change in m6A modification at 3’UTR of mRNAs tightly associates with the expression level of salt responsive genes, where show four regulatory mode including hyper-up, hyper-down, hypo-up and hypo-down, triangles indicate m6A modification, squares indicate gene expression, red and green color indicate up and down-regulation, respectively. The mRNAs with increased m6A level (dominantly enriched in photosynthesis pathway) are transported to the cytoplasm and recognized by m6A readers, which arises mRNA decay of these genes, resulting in a pronounced reduction of its transcript abundance. In tum, the changed photosynthesis capacity allows salt sensitivity or proper salt response in plants.

m^6^A-modified genes show significant enrichment in photosynthesis pathways, with more than 60% of m^6^A modifications localized near start codon of chloroplast-associated genes. Abundant m^6^A modification sites in photosynthetic genes suggest m^6^A may have specialized role in photosynthesis pathway. To determine whether m^6^A affects photosynthetic responses to salt stress, we measured Fv/Fm and NP Q using FluorCam in salt-treated and untreated *vir* mutants. While wild-type and *vir* mutants showed comparable Fv/Fm and NPQ under control conditions, salt stress reduced Fv/Fm significantly more and enhance NPQ significantly more in *vir* mutants (Fig 7C-F), indicating that VIR deficiency exacerbates photosystem II damage.

Given this m^6^A-dependent photosynthetic impairment, we examined chloroplast ultrastructure by TEM in *vir* and *ect2/3/4* mutants. Under control conditions, all genotypes exhibited intact chloroplasts with organized grana, though *vir* mutants displayed enlarged starch granules, suggesting disrupted starch metabolism. Under salt stress, the chloroplasts of wild-type and *ect2/3/4* mutants exhibited semicircular chloroplast curvature, reduced grana stacking, and disordered arrangement of thylakoid membranes (displaying a wavy configuration), accompanied by partial dissociation of the stromal lamellae and an increase in the number of starch granules and plastid globules. Strikingly, *vir* mutants showed fragmented chloroplasts with abnormal starch granules and complete loss of normal architecture under salt stress (Fig. 7g, h). These results demonstrate that VIR-mediated m^6^A modification maintains chloroplast integrity under salt stress, confirming its essential role in plant photosynthetic adaptation.

## Discussion

This study establishes that m^6^A epitranscriptomic reprogramming serves as a critical mechanism for plant salt stress tolerance. Our study demonstrates that salt stress triggers extensive transcriptional activation of Arabidopsis m^6^A regulatory machinery, including writers (*MTA*, *MTB*, *VIR*, *FIP37*, *HAKAI*), erasers (*ALKBH9B*/*10B*), and readers (*ECT2*/*3*/*4*), accompanied by a 15% increase in global m^6^A levels (Fig 1). This coordinated response signifies a fundamental role for epitranscriptomic reprogramming in salt adaptation. This aligns with previous reports linking m^6^A dynamics to abiotic stress responses. For instance, Arabidopsis m^6^A writer mutants (*mta*, *vir*) exhibit hypersensitivity to heat stress (Wei et al., 2018), while rice OsFIP37 knockdown impairs drought tolerance (Zhang et al., 2019). Our phenotypic analyses extend these observations, revealing that writer-deficient mutants (*mtb*, *vir*, *fip37*, *hakai*) display severe salt sensitivity (Fig 2), characterized by root growth inhibition, ROS accumulation, and biomass reduction, these phenotypes attributable to genotype by environment interactions rather than developmental defects, demonstrates that intact m^6^A methylation is essential for physiological and plant architecture maintenance under salt stress. Notably, the functional redundancy among readers ECT2/3/4 (Cai et al., 2024) was overcome by triple-mutant analysis, revealing their collective necessity for salt tolerance (Fig 3), while eraser mutants showed minimal phenotypes, likely due to compensatory homologs (Wei et al., 2018). These genetic analyses collectively position m^6^A machinery as a master regulatory switch in stress response pathways.

Despite global m^6^A hypermethylation, MeRIP-seq revealed that salt stress reshapes the landscape rather than the locus preference of m^6^A deposition, with enhanced enrichment in 3’-UTRs near stop codons (Fig 4B-E) and elevated signal intensity at transcription termination sites (TES) (Fig 4F). This suggests stress-specific recruitment of m^6^A machinery to 3’-regulatory zones, potentially fine-tuning mRNA stability, localization, or translation efficiency. This spatial redistribution mirrors findings in mammalian systems where stress-triggered m^6^A repositioning regulates mRNA stability (Zhou et al., 2015). The conserved UGUAH motif (Fig 4G) further implies sequence-directed specificity in stress-induced methylation, consistent with motifs reported by Shen et al. (2016). Crucially, 840 salt-responsive genes exhibited differential m^6^A modifications, with predominately negative correlation between gene expression and m^6^A level (Fig 5C-D), highlighting m^6^A-mediated decay as a major regulatory axis. Functional enrichment confirmed that >60% of differentially methylated genes under salt stress encode photosystem components, specifically PSII, Calvin cycle, and chloroplast ATPase components as a key target of m^6^A hypermethylation under salt stress (Fig 5E). Salt-induced hypermethylation accelerates decay of photosynthesis-related transcripts (e.g., *CA1*, *PNSB1*, *LHCB2.4*) via VIR-dependent m^6^A deposition (Fig 6). This negative regulation is consistent with the canonical role of m^6^A readers (e.g., YTHDF2) in promoting mRNA decay (Wang et al., 2014). It has been demonstrated that m^6^A demethylation components enhance the stability of the associated genes (Zhou et al., 2019). Research has shown that the C-terminal domain of YTHDF2 in the cytoplasm selectively binds to m^6^A-modified mRNA, while the N-terminal domain is responsible for directing YTHDF2-mRNA complexes to cellular RNA degradation sites, thereby accelerating the degradation of m^6^A-containing RNA. Six core photosynthesis genes showed elevated m^6^A in 3’-UTRs and reduced expression (Fig 6A-L) which accelerated decay in WT versus *vir* mutants (Fig 6M-R). Paradoxically, while suppressing photosynthesis may conserve energy under stress, complete loss of m^6^A-mediated control (*vir* mutant) exacerbated photodamage (reduced Fv/Fm, elevated NPQ; Fig 7C-F) and caused chloroplast disintegration (Fig 7G-H). This reveals a physiological trade-off, m^6^A-directed decay of photosynthetic transcripts prevents harmful protein production during stress, but requires precise tuning to avoid catastrophic energy deficit. Salt sensitivity of *ca1*, *pnsb1*, and *lhcb2.4* mutants (Fig 7A-B) confirms that controlled suppression of these genes is adaptive, whereas their unrestrained expression in *vir* disrupts organellar integrity. This finding extends earlier reports of m^6^A’s involvement in plastid RNA metabolism (Arribas-Hernández et al., 2018), highlighting its function beyond nuclear mRNA regulation. Our data support a model where salt-induced m^6^A hypermethylation acts as a transcriptional buffer and a critical architect of salt stress responses by predominately targeting photosynthetic genes, which will contribute new insights into the regulatory network of m^6^A modifications upon salt stress.

## Supplementary data

The following supplementary data are available at JXB online

Table S1. Primers used in this study for qPCR analyses.

Table S2. Quality control results of MeRIP-seq alignment.

Table S3. Differentially expressed genes detected in Col-0 response to salt stress.

Table S4. Differentially m6A peak detected in Col-0 response to salt stress.

Table S5. Correlation between differential m6A modiification and differential gene expression.

Fig S1. The expression of genes associated with m^6^A modification increases with the increase of salt treatment.

Fig S2. The effect of salt stress on the growth of the mutant m^6^A writer.

Fig S3. Effect of salt stress on growth of mutant m^6^A erasers *alkbh9b*, *alkbh10b* and *alknh9b/10b*.

Fig S4. Phenotypic analysis of mutants m^6^A reader.

Fig S5. T-DNA insertion sites related to photosynthesis mutants

Fig S6. Conduct a factor analysis of genotype-environment interactions to examine the G×E effects of *ca1*, *lhcb2.4*, *pnsb1* mutants under salt stress.

## Author contributions

Qinghua Yang, Data curation, Formal analysis, Validation, Investigation, Writing - original draft; Shuang Wang, Data curation, Validation, Investigation, Writing - original draft; Jieying Si, Validation; Qiuying Pang, Supervision, Funding acquisition, Project administration; Aiqin Zhang, Supervision, Funding acquisition, Writing - original draft, Project administration, Writing - review and editing.

Qinghua Yang and Shuang Wang contributed equally to this work.

